# Genome-wide screen of genomic imprinting in endosperm and population-level analysis reveal allelic variation for imprinting in flax

**DOI:** 10.1101/2020.11.20.390799

**Authors:** Haixia Jiang, Dongliang Guo, Jiali Ye, Yanfang Gao, Huiqing Liu, Yue Wang, Min Xue, Qingcheng Yan, Jiaxun Chen, Lepeng Duan, Gongzhe Li, Xiao Li, Liqiong Xie

## Abstract

Genomic imprinting is an epigenetic phenomenon caused by the biased expression of maternally and paternally inherited alleles. In flowering plants, genomic imprinting predominantly occurs in triploid endosperm and plays a vital role in seed development. In this study, we identified 241 candidate imprinted genes including 143 maternally expressed imprinted genes (MEGs) and 98 paternally expressed imprinted genes (PEGs) in flax (*Linum usitatissimum* L.) endosperm using deep RNA sequencing. The conservation of imprinting in plants is very limited and imprinting clustering is not a general feature. MEGs tends to be endosperm expression specific, while PEGs are non-tissue specific. Imprinted SNPs differentiated 200 flax cultivars into oil flax, oil-fiber dual purpose flax (OF) and fiber flax subgroups, suggesting that genomic imprinting contributes to intraspecific variation in flax. The nucleotide diversity (*π*) of imprinted genes in oil flax subgroup is significantly higher than that in fiber flax subgroup, indicating that some imprinted genes undergo positive selection during flax domestication from oil flax to fiber flax. Imprinted genes undergo positive selection is related to the functions. Eleven imprinted genes related to seed size and weight were identified using the candidate gene-based association study. Our study provides information for further exploring the function and genomic variation of imprinted genes in flax population.

## Introduction

Genomic imprinting is an epigenetic phenomenon occurring in both mammals and flowering plants (Hutter *et al*., 2010; Waters *et al*., 2013). Imprinting occurs in the placenta and embryo as well as in adult tissues in mammals (Renfree *et al*., 2012), while in flowering plants, imprinting predominantly occurs in endosperm, and rarely in embryo and seed coat (Yuan *et al*., 2017; Meng *et al*., 2018). The diploid embryo transmits genetic information to the next generation, while the triploid endosperm, with 2:1 ratio of maternal to paternal genomes (2m:1p), provides nutrition and signals for embryo development (Sabelli and Larkins, 2009; Mei *et al*., 2015). The deviation of the 2m:1p ratio in endosperm has adverse effect on seed development (Wang *et al*., 2018; Scott *et al*., 1998; Lu *et al*., 2012; Sekine *et al*., 2013), implying the important role of imprinting in endosperm development. Imprinted genes were studied only in a limited plant species, including monocots of rice (*Oryza sativa*) (Yuan *et al*., 2017; Luo *et al*., 2011), maize (*Zea mays*) (Waters *et al*., 2013; Meng *et al*., 2018; Zhang *et al*., 2011; Waters *et al*., 2011; Dong *et al*., 2017), sorghum (*Sorghum bicolor* L. Moench) (Zhang *et al*., 2016a) and dicots of *Arabidopsis* (Gehring *et al*., 2011; Wolff *et al*., 2011; Hsieh *et al*., 2011), castor bean (*Ricinus communis*) (Xu *et al*., 2014). In most dicots, the endosperm is transient and consumed by the embryo at the later stage of seed development (Sreenivasulu and Wobus, 2013), while the endosperm of most monocots remains persistently and serves as a source of nutrition for seed germination (Luo *et al*., 2011). There is no extensive conservation of imprinted genes across species (Waters *et al*., 2011; Zhang *et al*., 2016a; Dong, 2017), suggesting different species may require a unique set of imprinted genes for seed development.

In plants, imprinted genes were detected to be involved in the regulation of seed development, seed dormancy and postzygotic reproductive isolation (Sun *et al*., 2017; Piskurewicz *et al*., 2016; Kradolfer *et al*., 2013; Wolff *et al*., 2015). The loss of function of some imprinted genes leads to seed abortion (Joanis and Lloyd, 2002; Chaudhury *et al*., 1997; Berger *et al*., 2006). In *Arabidopsis*, the maternally expressed imprinted gene *MEA* encodes a Polycomb Repressive Complex 2 (PRC2) subunit in Polycomb group (PcG) complex. The seeds with maternal *MEA* allele developed normally, while those carrying maternal *mea* allele were aborted, regardless of the genotypes of paternal allele (Joanis and Lloyd, 2002). Imprinted genes influence seed size by regulating the development of embryo and endosperm (Yuan *et al*., 2017; Scott *et al*., 1998; Köhler *et al*., 2005; Chen *et al*., 2016). In rice, the loss-of-function mutants of *MEG2* and *MEG3* (MEGs) showed a significant reduction in seed size and weight, and the loss-of-function of *PEG1, PEG2, PEG3* (PEG) decreased starch content, seed size and yield (Yuan *et al*., 2017). Regarding to another imprinted gene *OsFIE1* in rice, RNAi lines and homozygous T-DNA insertion mutant *osfie1* lines all showed delayed embryo development and reduction of seeds fertility, grain size, grain weight and aleurone layer cells (Huang *et al*., 2016). Even a few imprinted genes were shown to be important in seed development, there are many more, of which the function is yet to be determined.

Some imprinted genes were shown to be under positive selection and intraspecific variation features (Hutter *et al*., 2010; Berger *et al*., 2012; Pignatta *et al*., 2014). *MEA* originated a block duplication 35 to 85 million years ago owing to a whole-genome duplication within the Brassicaceae lineage (Spillane *et al*., 2007). After duplication, *MEA* underwent positive selection consistent with neo-functionalization and the parental conflict theory (Spillane *et al*., 2007; Miyake *et al*., 2009). In maize, conservative imprinting genes increased the substitution rate of nonsynonymous to synonymous (dN/dS) compared with non-conservative imprinting genes and more likely to undergo positive selection (Waters *et al*., 2013). PEGs were more likely to be under positive selection and rapidly evolve than MEGs in *Arabidopsis thaliana* (Tuteja *et al*., 2019). Imprinting showed evidence of intraspecific variation in *Arabidopsis* and maize (Waters *et al*., 2013; Pignatta *et al*., 2014). The existence of intraspecific variation of imprinting was associated with epigenetic variation (Pignatta *et al*., 2014).

Flax (*Linum usitatissimum* L.) is an important economic crop due to its stem fiber and seed oil (Cloutier *et al*., 2012; Guo *et al*., 2020). Flax is a strict annual self pollination crop with a smaller genome size (∼373 Mb) (Wang *et al*., 2012), which are good for biological research. Cultivated flax were domesticated from a pale flax (*Linum bienne*) for oil usage at 10,000 years ago in the Near East and differentiated into fiber, OF and oil subgroups during the domestication of flax (Guo *et al*., 2020; Allaby *et al*., 2005; Fu and Allaby, 2010). Some genes underlying important traits including plant architecture, flowering, dehiscence, oil production and yield underwent strong artificial selection in the domestication process of flax (Guo *et al*., 2020; Zhang *et al*., 2020). Whether imprinted genes are also undergone artificial selection during flax domestication has not been reported.

In this study, we have performed RNA-seq analysis of flax endosperm isolated from the reciprocal crosses between CIli2719 and Z11637. Based on the parent-of-origin biased expression of the parental alleles in the endosperm from both crosses, we identified 241 moderately imprinted genes including 143 MEGs and 98 PEGs, 67 strongly imprinted genes including 63 MEGs and 4 PEGs and 19 completely MEGs. The analysis of imprinted genes at population level demonstrated that imprinted genes divided the 200 flax germplasms into oil flax, OF and fiber flax subgroups, which suggested that imprinting promoted intraspecific variation of flax. The nucleotide diversity analysis showed that some imprinted genes were shown to be under positive selection consistent with function. Furthermore, we identified 11 imprinted genes associated to seed size and weight. Our results will provide a theoretical basis for further study of gene imprinting and provide some insights for understanding the diversity of imprinted genes.

## Results

### Identification of imprinted genes in flax endosperm

To understand the parental origin of gene expression in flax, we performed RNA-seq analysis in endosperm isolated from the F_1_ generation of CIli2719×Z11637 (CZ) and Z11637×CIli2719 (ZC) at 7 days after pollination (DAP). Average 7.21 Gb of 125bp paired-end clean reads were obtained from each biological replicate with the Illumina novaseq6000 platform. The clean reads from CZ and ZC were aligned to parental genomes CIli2719 and Z11637 (Guo *et al*., 2020) to identify the reads specifically originated from one of parents at each SNP site for allelic expression (Figure S1). In total, 52,794 SNPs in both reciprocal crosses with at least ten reads could be assigned to a parental allele and were used for allele-specific expression analysis in hybrid endosperm. About 37,000 SNP loci showed statistically significant deviation (*p*<0.05, χ^2^ test) from the expected 2m:1p ratio in both CZ and ZC endosperm.

Three different thresholds were used to identify genes showing parent-of-origin biased expression at three different levels (see “Materials and methods”). Among these SNP loci, 498 loci were considered as moderately imprinted loci, including 319 maternally expressed imprinted SNP loci (ME-SNPs) corresponding to 143 MEGs and 179 paternally expressed imprinted SNP loci (PE-SNPs) corresponding to 98 PEGs. And 141 loci were identified as strongly imprinted loci, including 135 ME-SNPs and 6 PE-SNPs which correspond to 63 MEGs and 4 PEGs, respectively. In addition, 36 loci were identified as completely imprinted loci and all of them were ME-SNPs corresponding to 19 MEGs (Figure 1, Table S1). Among 241 imprinted genes, 229 genes were protein-coding genes (Table S2).

**Figure 1.**
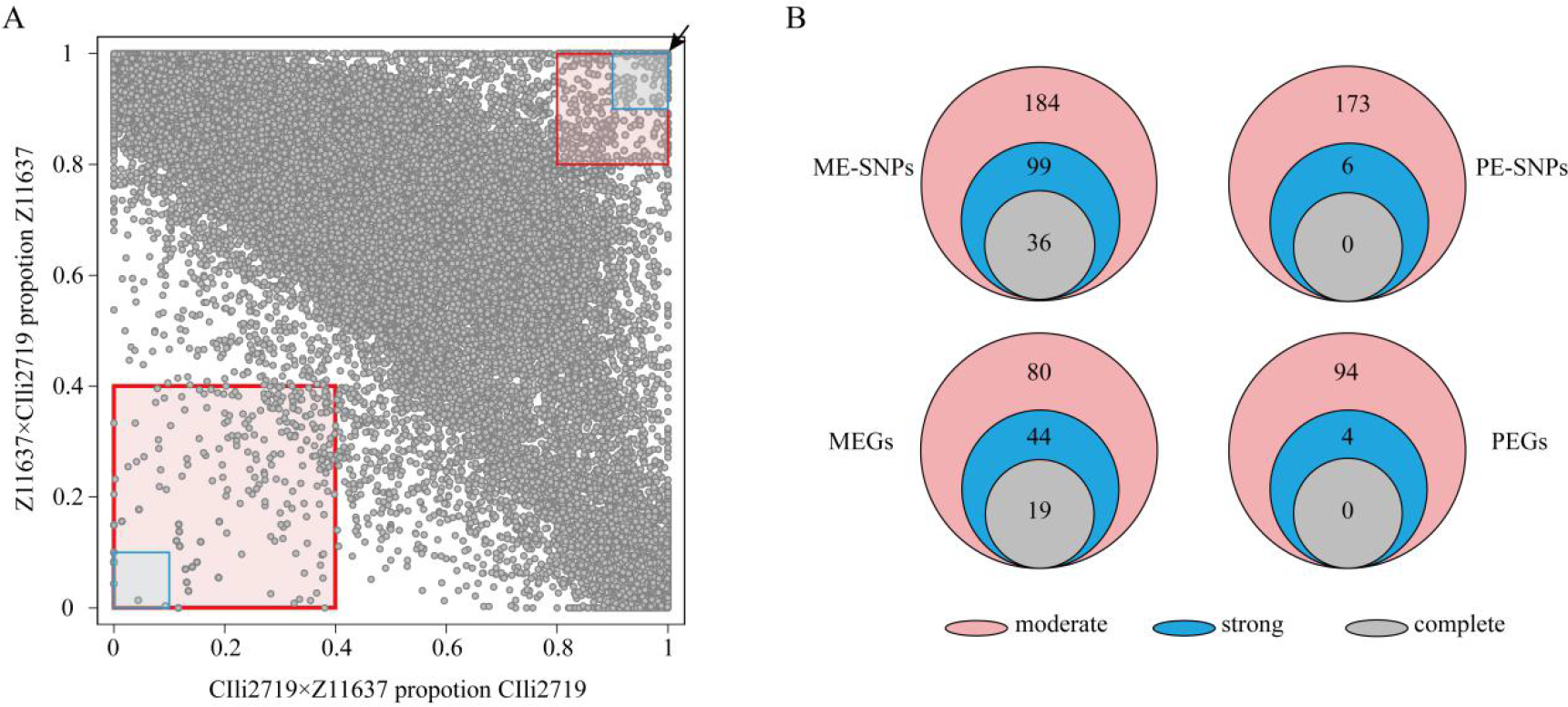
Identification of imprinted genes in flax endosperm. (A) The proportion of parental transcripts in both CZ and ZC was plotted for 52,794 SNPs with at least ten reads could be assigned to a specific allele. The shaded areas indicated moderate (pink), strong (blue), or complete (arrows) maternally expressed imprinted SNP loci (upper right) or paternally expressed imprinted SNP loci (lower left). (B) The number of moderate (pink), strong (blue), and complete (gray) imprinted SNP loci and imprinted genes in endosperm. ME-SNPs, maternally expressed imprinted SNP loci; PE-SNPs, paternally expressed imprinted SNP loci; MEG, maternally expressed imprinted genes; PEG, paternally expressed imprinted gene.

### Validation of imprinted genes

For experimental confirmation, thirteen genes including five MEGs, five PEGs, and three non-imprinted genes, which represented the whole transcripts by RNA sequencing in this study, were randomly selected to validate the gene expression level of the high throughput sequencing with qRT-PCR analysis. The analysis showed that the gene expression level of the selected genes by qRT-PCR analysis was consistent with the RNA-seq data (Figure S2).

For further verification of the imprinting status, nine MEGs and three PEGs were used to perform RT-PCR on the hybrid endosperm and parents endosperm followed by Sanger sequencing (Figure S3). A 400-800bp fragment of each gene with at least one imprinted SNP site was selected for PCR amplification. The results showed that nine MEGs were predominantly expressed from the maternal alleles and three PEGs were preferentially expressed from the paternal alleles in reciprocal crosses, which were consistent with the RNA-seq data.

### Characterization analysis of imprinted genes identified in flax

We carried out gene ontology (GO) analysis for the 229 imprinted protein-coding genes of flax, including 135 MEGs and 94 PEGs (Table S2). The background genes for GO analysis were 12,395 endosperm-expressed genes with at least ten reads could be assigned to a specific allele in both CZ and ZC. Categories with a significant level (*P*<0.05) were defined as enriched. Compared with the whole transcripts in endosperm, imprinted genes were significantly enriched in catalytic activity or transferase activity according to their molecular function and metabolic process according to their biological processes (Figure S4A, Table S3).

To evaluate the interspecific conservation of imprinted genes, we compared imprinted genes identified in this study with the imprinted genes reported in *Arabidopsis* (Gehring *et al*., 2011), rice (Yuan *et al*., 2017; Luo *et al*., 2011), maize (Waters *et al*., 2013; Dong, 2017), sorghum (Zhang *et al*., 2016a), and castor bean (Xu *et al*., 2014). The analysis showed that there were 115, 89 and 59 genes found to be homologous in at least one of the five species at different confidence levels (<1E-10, <1E-20 and <1E-50, respectively) (Figure 2, Table S4-S5). At E-value<1E-10, there were 32, 26, 43, 52, 39 flax imprinted genes conserved in *Arabidopsis*, castor bean, rice, sorghum and maize, respectively (Figure 2A, Table S4-S5). And 19, 21, 35, 46, 27 imprinted genes conserved with these plants at E-value<1E-20, respectively (Figure 2B, Table S4-S5). In addition, less imprinted genes 13, 11, 21, 22 and 15 conserved with the five species at E-value<1E-50, respectively (Figure 2C, Table S4-S5). Some imprinted genes had imprinted homologs in up to four species, while no imprinted gene in flax was conserved in all species (Figure 2, Table S4-S5). These results suggested that the conservation of imprinting in plants was quite limited.

**Figure 2.**
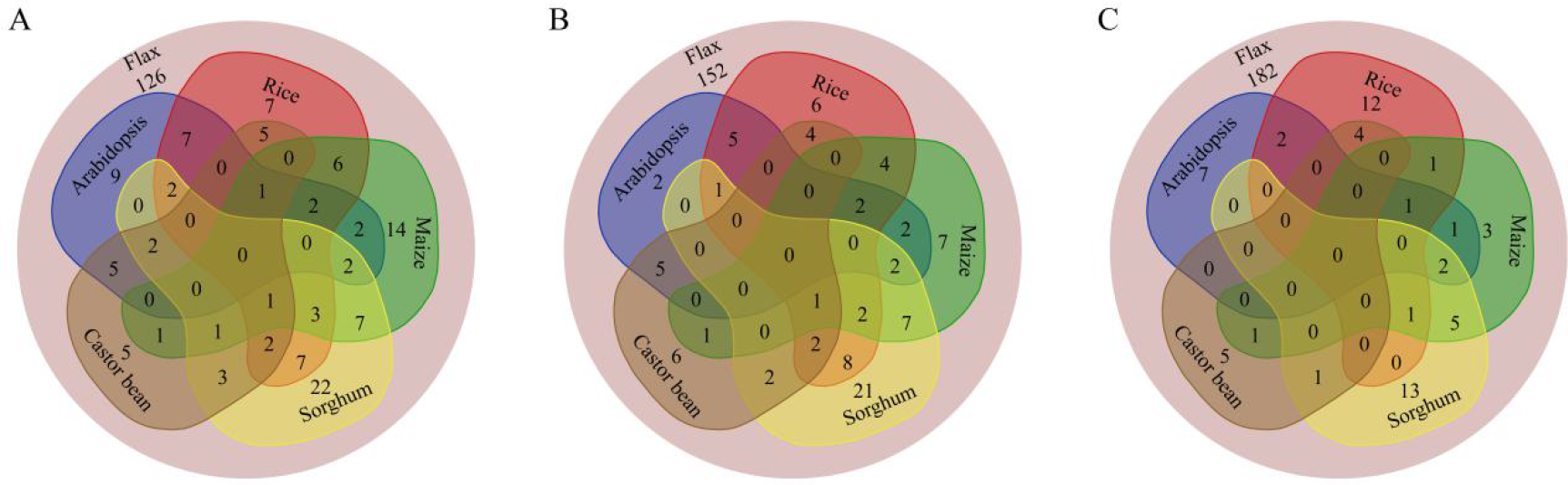
The conservation of flax imprinted genes between other species. (A) Venn diagram showing overlaps of imprinted genes at E-value<1E-10 between flax and maize, rice, *Arabidopsis*, castor bean, sorghum, respectively. (B) Venn diagram showing overlaps of imprinted genes at E-value<1E-20. (C) Venn diagram showing overlaps of imprinted genes at E-value<1E-50.

Intriguingly, the expression of some conserved flax imprinted genes showed different parental origin in other species. For example, the flax gene *Lus10022747* encoding a serine/threonine-protein kinase WNK5 and its homologues in *Arabidopsis*, castor bean and rice showed maternally preferential expression, while its maize homolog displayed preferentially paternal expression (Table S4). A PAS domain tyrosine kinase family protein-coding gene *Lus10040540* and its homolog of maize were PEGs, but its homologues in castor bean, rice and sorghum were MEGs (Table S4). Among 115 conserved flax imprinted genes, only 59% (58 MEGs and 10 PEGs) remain the same preference of parental expression with other species (Table S4).

GO enrichment analysis of 115 conserved imprinted genes in flax displayed that MEGs significantly enriched in catalytic activity and metabolic process according to their molecular function and biological processes, respectively. PEGs enriched in compound binding according to their molecular function (Figure S4B, Table S6).

### Clustering of the flax imprinted genes

To study the genomic distribution of flax imprinted genes, 229 imprinted genes were mapped to fifteen chromosomes for cluster analysis. The 229 imprinted genes were scattered distribution across fifteen chromosomes. By analyzing the genomic distance between the imprinted genes, we found that most of them were not co-localized in a cluster, and only 24 were fall into 12 clusters, where two imprinted genes of each cluster were within 10 kb (Figure 3, Table S7). The finding was similar to the results of *Arabidopsis* (Gehring *et al*., 2011; Wolff *et al*., 2011), maize (Waters *et al*., 2011), rice (Luo *et al*., 2011), castor bean (Xu *et al*., 2014) and sorghum (Zhang *et al*., 2016a), showing that clustering of imprinted gene is not a common phenomenon in plants.

**Figure 3.**
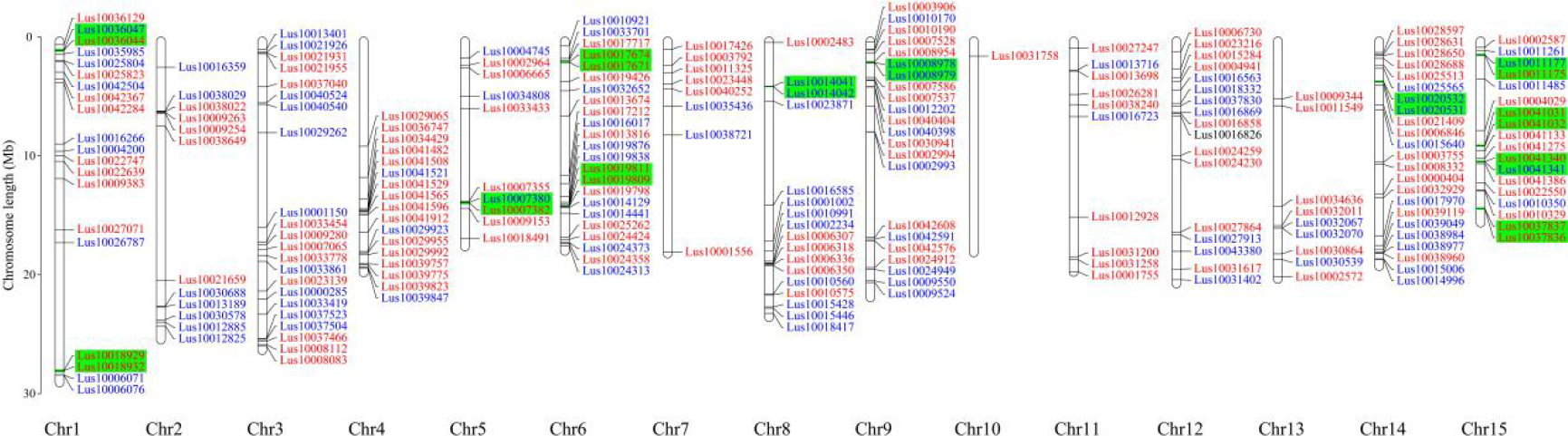
Some flax imprinted genes are located in mini-clusters. The MEGs (red) and PEGs (blue) were mapped to the 15 flax chromosomes. Genes clustered within 10 kb are boxed in green.

### Endosperm-specific expression of the flax imprinted genes

We analyzed the expression specificity of the imprinted genes in various tissues. The majority of MEGs (60%) preferentially expressed in endosperm, and only a minority (25.5%) of PEGs were preferentially expressing in endosperm (Figure 4A-B). The expression level of endosperm-preferred MEGs (endo-MEGs) and PEGs (endo-PEGs) were significantly (*P*<0.05) higher than that of all genes, whereas there was no evidence that endo-MEGs or endo-PEGs exhibited unusually high or low expression levels than other MEGs or PEGs which also expressed in other tissues (Figure 4C-D). We also analyzed the tissue specificity of 115 (75 MEGs and 40 PEGs) conserved imprinted genes and 38 (36 MEGs and 2 PEGs) conserved strong imprinted genes. Among 115 conserved imprinted genes, 49 (65.3%) MEGs and 12 (30%) PEGs showed endosperm-preferred expression (Figure S5A-B), and among the 38 conserved strong imprinted genes, 29 MEGs (80.6%) and all PEGs were preferentially expressed in endosperm (Figure S5C), respectively. These results suggested that MEGs and the conserved imprinted genes are more likely to be preferentially expressed in endosperm.

**Figure 4.**
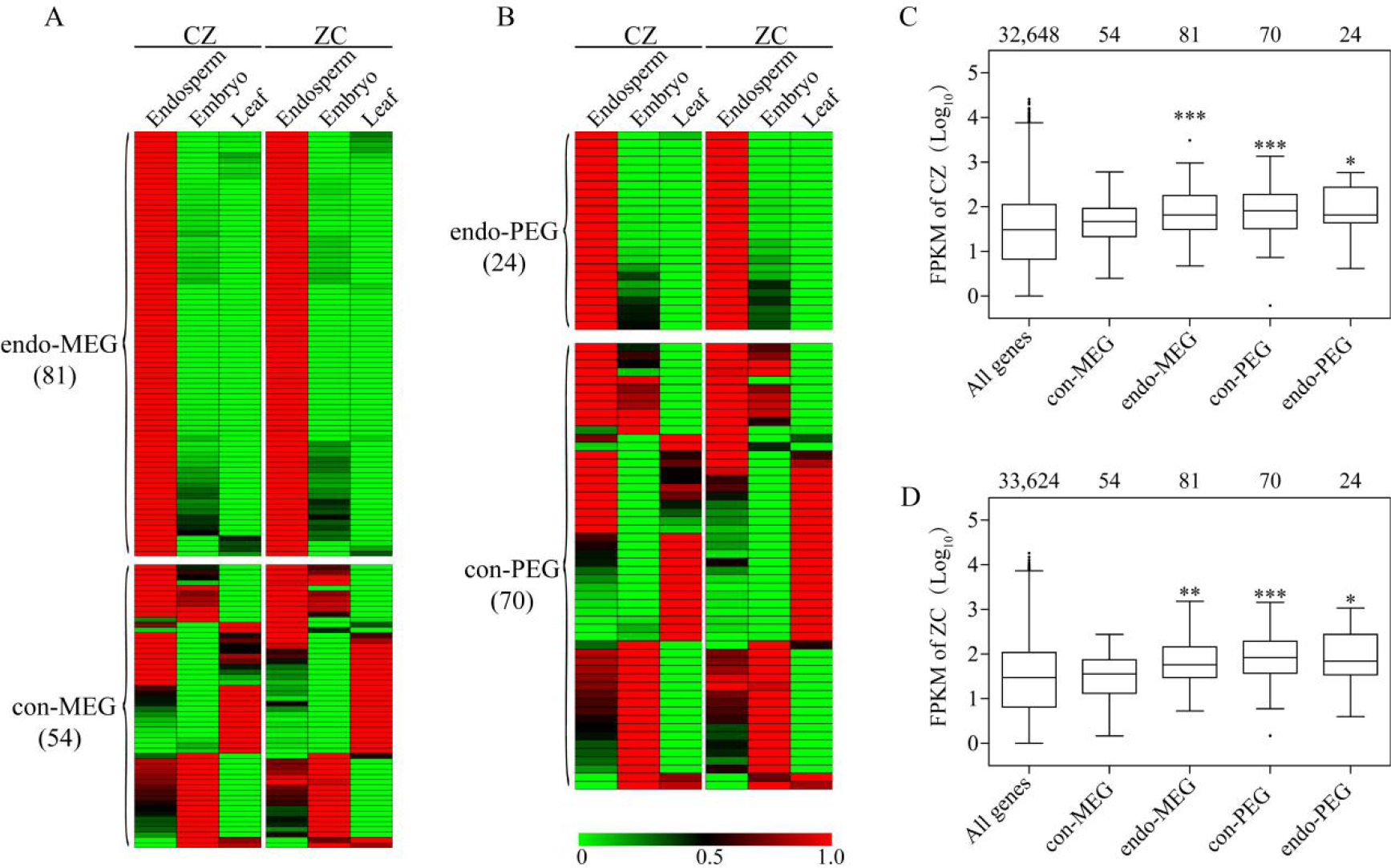
Expression of flax imprinted genes over various flax tissues in both reciprocal hybrids based on RNA-seq analysis. (A, B) The gene-expression patterns for MEGs (A) and PEGs (B). endo-MEGs, MEGs that expressed preferentially in endosperm; con-MEGs, MEGs that also expressed in other tissues; endo-PEGs, PEGs that expressed preferentially in endosperm; con-PEGs, PEGs that also expressed in other tissues. The normalized values were used for hierarchical clustering and the heat map indicated relative levels of expression. The endosperm and embryo tissues were harvested at 7 DAP and the leaf tissues were collected at 2 weeks after planting. For each sample, three biological replicates were used. (C, D) The Log_10_ (FPKM) values of CZ (C) and ZC (D). The Log_10_ (FPKM) values of CZ (C) and ZC (D). All genes with FPKM>1 in endosperm were used in this study. The values listed above box plots were the number of genes in each group. Asterisks indicate the significance level (*, *P* < 0.05; **, *P* < 0.01; ***, *P* < 0.001).

### Flax imprinted genes can differentiate flax subgroups

To investigate whether the variation in imprinted genes reflects genetic diversity among 200 natural flax varieties, we detected individual kinship of imprinted SNPs (498) or genome-wide SNPs (674,074) (Guo *et al*., 2020). The kinships between imprinted SNPs or genome-wide SNPs were significantly correlated (*R*^2^=0.8457) (Figure 5A). Phylogenetic tree constructed based on all imprinted SNPs, ME-SNPs or PE-SNPs separated 200 accessions into three different subgroups which correspond to oil flax, OF and fiber flax subgroups (Figure 5B, Figure S6). The flax population could also be separated into three subgroups by principle component analysis (PCA) (Figure 5C). These results indicated that the allele frequency of imprinted SNPs was significantly different among subgroups, suggesting that imprinted genes may be selected differently in subgroups and contribute to domestication.

**Figure 5.**
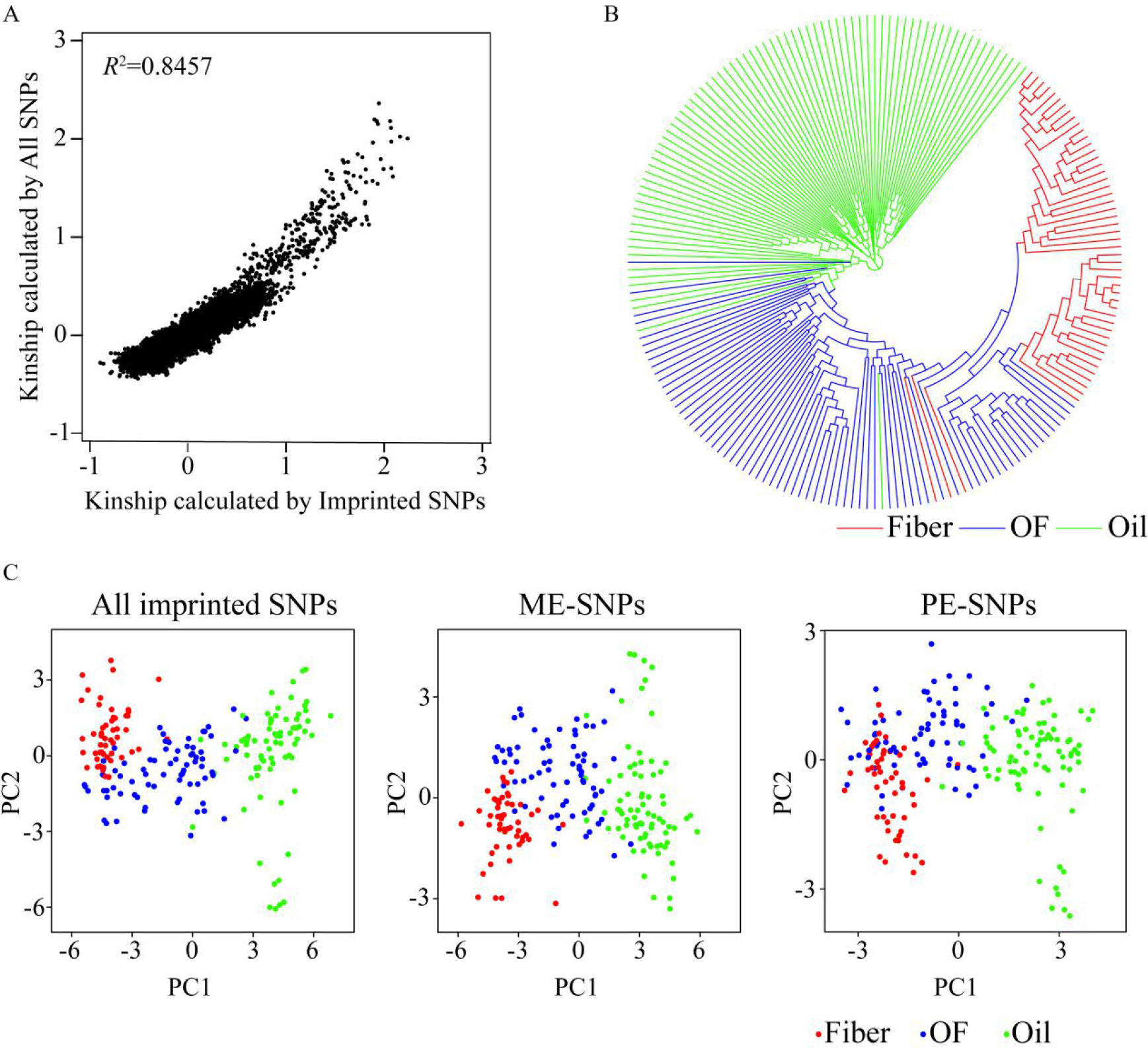
Imprinted genes differentiate flax subgroups. (A) The correlation of kinships between imprinted SNPs or all SNPs. (B) Phylogenetic tree of 200 flax accessions inferred from imprinted SNPs. (C) PCA plots of all imprinted SNPs, ME-SNPs and PE-SNPs. Fiber flax, oil-fiber dual purpose flax (OF), and Oil flax were represented in red, blue and green colors, respectively.

### Selective sweep signals in imprinted genes

To test the hypothesis that the diversity of imprinted genes are different in flax population, we collated all SNPs (3,191 SNPs) of 241 imprinted genes and compared the nucleotide diversity between oil flax (78 germplasms) and fiber flax subgroups (51 germplasms) which represented two primary morphotypes of cultivated flax (Zhang *et al*., 2020). The π values of all imprinted genes, MEGs or PEGs decreased significantly in fiber flax compared with oil flax (*P* < 0.0001, *t* test) (Figure 6A-C). We focused on two imprinted genes, *Lus10010350* (PEG) and *Lus10024230* (MEG), which contained more SNPs (31 SNPs and 43 SNPs, respectively) in genomic sequences. The allele frequency distribution of imprinted SNPs in *Lus10010350* was significantly different in the two subgroups, and the alternate allele ‘G’ (position 333406) was primarily found in the oil subgroup and rarely in fiber subgroup (Figure 6D). Significant reduction of π was also observed at the *Lus10010350* locus in fiber subgroup compared with that of oil subgroup and 90.32% of the SNPs were identified with a signature of purifying selection (Figure 6E-F). Similarly, the allele frequency distribution of *Lus10024230* was obvious different between the two subgroups and the alternate allele ‘A’ (position 492363) accounted for 61.67% in oil subgroup but only 6.98% in fiber subgroup (Figure 6G). Compared with oil subgroup, the π value of the SNPs in *Lus10024230* was dramatically decreased in fiber subgroup, and 95.35% of the SNPs were identified with a signature of purifying selection (Figure 6H-I). Taken together, these findings suggested that some imprinted genes may have been subjected to artificial selection during flax domestication.

**Figure 6.**
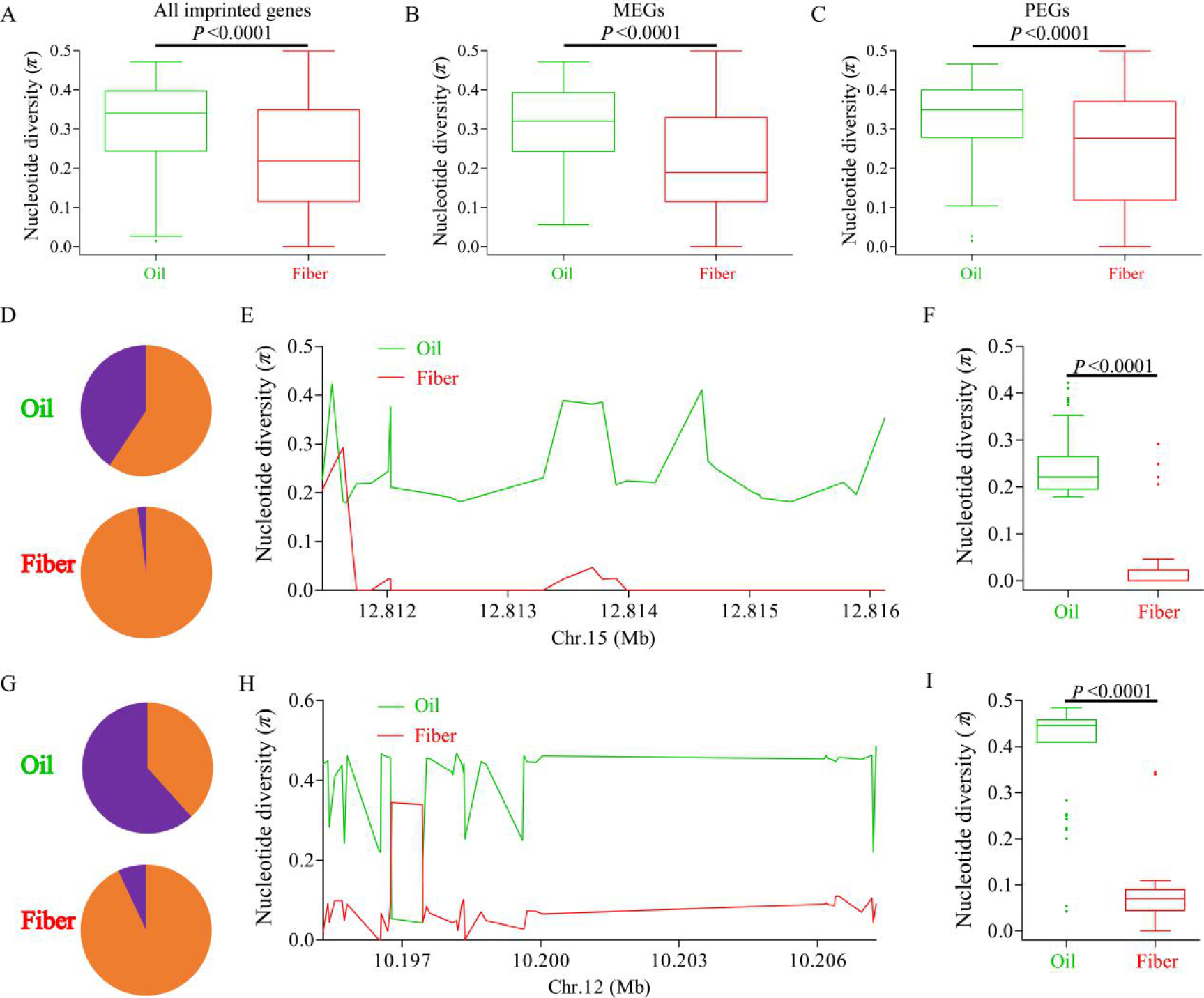
Distribution of nucleotide diversity (*π*) within imprinted genes and allele frequency differences of two genes across oil and fiber subgroups. (A-C) Boxplots for nucleotide diversity of all imprinted genes (A), MEGs (B) and PEGs (C) across Oil (green) and Fiber (red) groups. (D, G) The distribution of allele frequency of SNPs located in *Lus10010350* (D, PEG) and *Lus10024230* (G, MEG) in Oil and Fiber subgroups. The alternate alleles and reference alleles were shown in purple and orange, respectively. (E, H) The nucleotide diversity distribution of *Lus10010350* on chromosome 15 (E) and *Lus10024230* on chromosome 12 (H) among Oil and Fiber subgroups. (F, I) Boxplots for nucleotide diversity of *Lus10010350* (F) and *Lus10024230* (I) among Oil and Fiber subgroups. The difference was analyzed by two-tailed *t* tests.

### Candidate gene-based association study for seed size using flax imprinted genes

Imprinted genes played an important role in the regulation of endosperm development and seed size (Yuan *et al*., 2017; Luo *et al*., 2000; Guitton *et al*., 2004; Huang *et al*., 2017; Kinoshita *et al*., 1999; Kiyosue *et al*., 1999). To investigate whether imprinting is associated with seed size in flax, imprinted genes were used to perform candidate gene-based association study of seed size-related traits including seed length (SL), seed width (SW) and 1,000-seed weight (1000-SW). Using the general linear model (GLM) and mixed linear model (MLM) in TASSEL 5.0 (Bradbury *et al*., 2007), 33 imprinted genes containing 63 associated loci (SNPs) were detected to be associated with seed size and weight. Among them, 11 imprinted genes were repeatedly detected at least two environments or traits (Figure 7A-B, Table S8).

**Figure 7.**
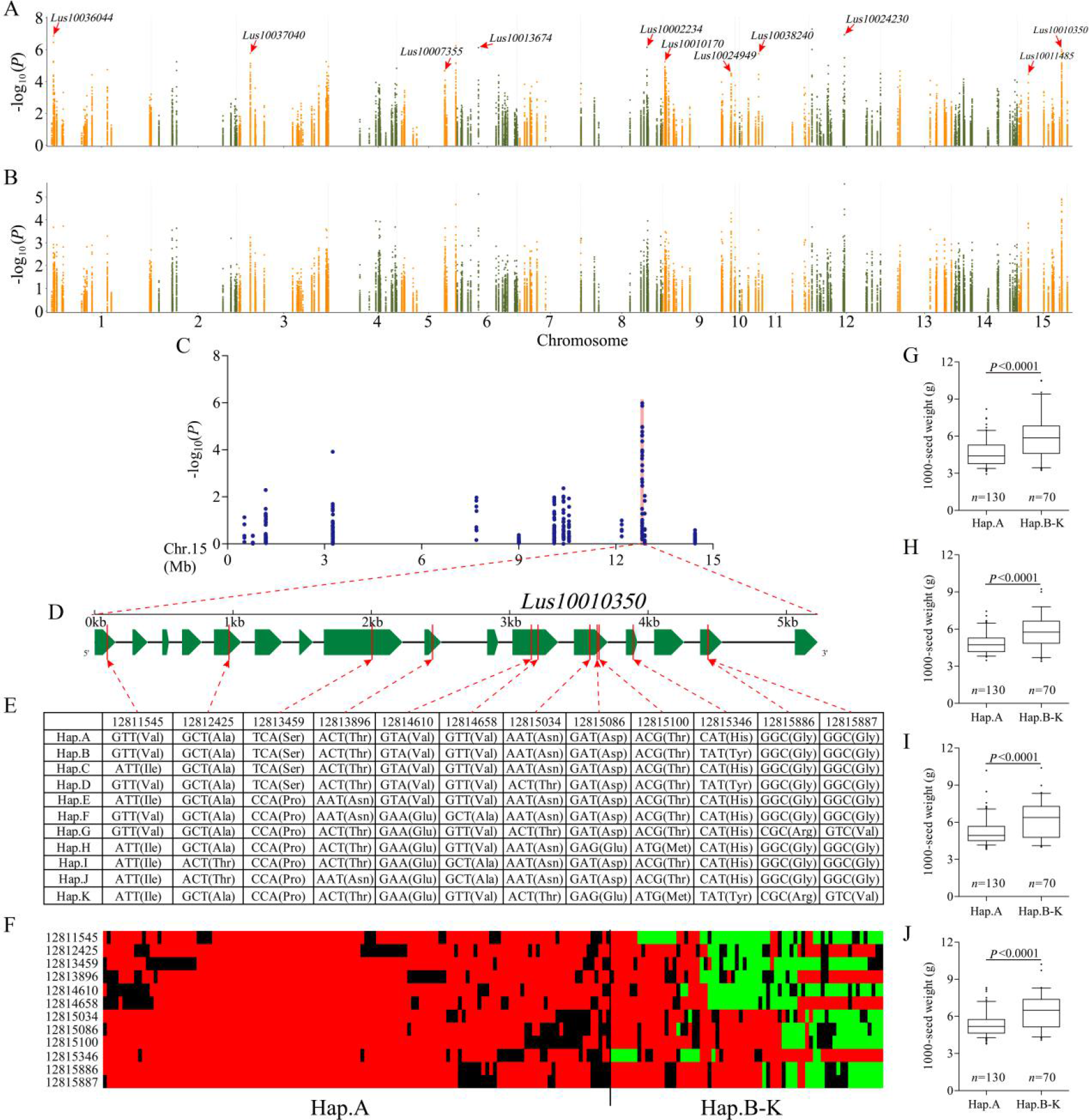
Imprinted gene-based association study for seed size and weight, and identification of a causal gene for the peak on chromosome 15. (A-B) The overlapping Manhattan plots for seed length, seed width and 1,000-seed weight in four environments (including 2016DL, 2017UR, 2019UR, 2019YL) using GLM (A) and MLM (B) models. Imprinted genes repeatedly detected at least two environments or traits related to seed size and weight were marked by red arrows. (C) Local Manhattan plots for seed width in 2016DL of imprinted gene-based association analysis surrounding the peak on chromosome 15. The position of *Lus10010350* was highlighted by a shaded pink column. (D) Gene structure of *Lus10010350*. (E) DNA polymorphism in *Lus10010350*. (F) Schematic representation of the structural variation in *Lus10010350*. (G-J) Boxplots for 1,000-seed weight based on the haplotypes (Hap.) for *Lus10010350* in 2016DL (G), 2017UR (H), 2019UR (I) and 2019YL (J). In the box plots, the center line represented the median, box limits indicated the upper and lower quartiles, whiskers marked the range of the data and points showed outliers. *n* indicated the number of accessions with the same genotype. The difference between haplotypes was analyzed by two-tailed *t* tests.

One of the significant signal peaks on chromosome 15 contained 9 repetitive SNPs (Figure 7C, Table S8), which located in the *Lus1001035*0 (PEG), encoding a bifunctional arginine demethylase and lysine hydroxylase jmjd6 protein. This gene contained 31 SNPs, of which 12 induced nonsynonymous mutations and formed 11 haplotypes (Figure 7D-E). We classified 200 accessions into two groups including haplotype A (reference alleles) and haplotypes B–K (alternate alleles) based on gene structural variation (Figure 7F). We found that flax accessions in haplotypes B–K had significantly longer seed length and width, and larger 1,000-seed weight than those in haplotype A (Figure 7G-J, Figure S7). These results suggested that the PEG *Lus10010350* may be involved in the seed size regulation in flax.

## Discussion

### Characterization of imprinted genes in flax endosperm

GO analysis for the 229 imprinted protein-coding genes of flax revealed that a majority of imprinted genes were significantly enriched in catalytic activity and metabolic process (Figure S4A, Table S3), similar to the results in castor bean (Xu *et al*., 2014) and sorghum (Zhang *et al*., 2016a), suggesting imprinted genes affected various aspects of endosperm development. However, imprinted genes have limited conservation across plant species, in contrast to those in mammals (Waters *et al*., 2011; Zhang *et al*., 2016a; Xu *et al*., 2014; Dong, 2017; Zhang *et al*., 2003). In maize and sorghum endosperm, only about 10% and 33% imprinted genes were conserved with other species, respectively (Waters *et al*., 2011; Zhang *et al*., 2016a). Among 165 imprinted genes in rice, only 33% of them were conserved in maize (Dong, 2017). Twenty-five (12%) imprinted genes identified in castor bean were conserved in *Arabidopsis*, rice or maize (Xu *et al*., 2014). In this study, the conservation of flax imprinted genes was evaluated with those in other species (*Arabidopsis*, castor bean, rice, sorghum and maize) at three levels of stringency. There were 115 (50.2%), 89 (38.9%), 59 (25.8%) imprinted genes in flax which were homologous to those in at least one of the five species at E-value<1E-10, 1E-20 and 1E-50, respectively (Figure 2, Table S4-S5) while none imprinted genes of flax were found having imprinted homologs with all five species (Figure 2, Table S4-S5). Those results suggested that some common pathways in different flowering plants may need to be regulated by imprinting to modulate endosperm development but different genes in the pathway are selected to be imprinted in different species. This explains why the conservation of imprinting in plants is quite limited.

An interesting observation was that among 115 conserved imprinted genes, only 68 genes (58 MEGs and 10 PEGs) showed the same origin of parental expression as in other species (Table S4). The remaining 47 genes (17 MEGs and 30 PEGs) had opposite origin of parental expression. For instance, the gene *Lus10041031* is a complete MEG identified in our study, but its maize homolog is a PEG (Waters *et al*., 2013). *Lus10021926* is a strong PEG in flax, while in *Arabidopsis* and castor bean its homologies are MEGs (Gehring *et al*., 2011; Xu *et al*., 2014). The homologous genes of the strong PEG *Lus10016563* are PEG in maize and MEGs in rice and sorghum (Waters *et al*., 2013; Yuan *et al*., 2017; Zhang *et al*., 2016a). This suggested that those genes have opposite mode of parental expression in different species may be subject to gene dosage, a mechanism thought to be important for endosperm development (Wang *et al*., 2018; Scott *et al*., 1998; Lu *et al*., 2012; Sekine *et al*., 2013). It was noteworthy to note that the gene *Lus10041386* encoding a histone-lysine N-methyltransferase Enhancer of Zeste homolog 2 (EZH2) is homologous to the FERTILIZATION-INDEPENDENT SEED Polycomb Repressive Complex 2 (FIS-PRC2) class gene *MEA* in *Arabidopsis* and the imprinted gene *GRMZM2G157820* (*EZH2*) in maize (Table S4). In rice and other species, other members of PRC2 genes are also imprinted, suggesting that imprinting of PRC2 genes is a conserved mechanism in flowering plants. Several lines of evidence suggested that PRC2 repressed the replication of central-cell nuclear before fertilization likely by the maternally expressed alleles and regulated endosperm proliferation, suggesting a vital role in seed development (Chaudhury *et al*., 1997; Kiyosue *et al*., 1999; Zhang *et al*., 2005; Ohad *et al*., 1996; Ohad *et al*., 1999; Luo *et al*., 1999; Moreno-Romero *et al*., 2019).

### Imprinted genes are not extensively clustered

Physical clustering of imprinted genes is a conserved feature in mammals (Gehring *et al*., 2011; Gregg *et al*., 2010), while there is little evidence of clustering in plant species. Imprinted genes identified in maize (Waters *et al*., 2011), *Arabidopsis* (Gehring *et al*., 2011; Wolff *et al*., 2011), rice (Luo *et al*., 2011), castor bean (Xu *et al*., 2014) and sorghum (Zhang *et al*., 2016a) were not shown to be extensive clustered. Using a clustering criterion consistent with that in *Arabidopsis* (∼125 Mb) and castor bean (∼350 Mb) which have comparable genome size to flax (∼373 Mb) (Gehring *et al*., 2011; Xu *et al*., 2014), we found that 24 of 229 flax imprinted genes were fall into 12 clusters (Figure 3, Table S7), similar to the proportion in *Arabidopsis* (Gehring *et al*., 2011; Wolff *et al*., 2011), rice (Luo *et al*., 2011), maize (Waters *et al*., 2011), castor bean (Xu *et al*., 2014) and sorghum (Zhang *et al*., 2016a), suggesting that imprinting clustering may be not a general feature in plants. Whether the clustered imprinted genes are coordinately regulated as those genes in animal clusters remains to be investigated.

### Endosperm-specific expression of flax imprinted genes

According to previous reports, imprinted genes in plants were mainly restricted to express in endosperm (Berger *et al*., 2012). But more and more studies had shown that only some imprinted genes are preferentially expressed in endosperm, while others are also expressed in other tissues (Waters *et al*., 2013; Waters *et al*., 2011; Dong, 2017). The proportions of endo-MEGs and endo-PEGs were dramatically different (68% MEGs versus 26% PEGs, 51% MEGs versus 24% PEGs in maize; 50% MEGs versus 16% PEGs in rice; 50% MEGs versus 20% PEGs in sorghum) (Waters *et al*., 2013; Dong, 2017). In our study, we found 81 endo-MEGs (60%) and only 24 endo-PEGs (25.5%) in flax (Figure 4A-B), similar to the proportion in rice, sorghum and maize (Waters *et al*., 2013; Zhang *et al*., 2016a; Dong, 2017). The expression level of endo-MEGs and endo-PEGs was significantly higher than that of all genes (Figure 4C-D). Compared with all imprinted genes (60% endo-MEGs versus 25.5% endo-PEGs), the proportion of endo-MEGs (65%) and endo-PEGs (30%) of conserved imprinted genes increased (Figure S5A-B). In the 38 (36 MEGs and 2 PEGs) conserved strong imprinted genes, 29 MEGs (80.6%) and all PEGs were endosperm-preferred expression (Figure S5C). These findings suggested that MEGs tends to be endosperm preferentially expressed, while PEGs are inclined to non-tissue specific expression. It also implied that the conserved imprinted genes are more likely to be preferentially expressed in endosperm and play an important role in seed development.

### Candidate gene-based association study reveals that some imprinted genes are involved in flax seed size regulation

Previous studies have shown that imprinted genes play an important role in seed development by regulating the development of endosperm (Yuan *et al*., 2017; Köhler *et al*., 2005; Chen *et al*., 2016; Luo *et al*., 2000; Guitton *et al*., 2004; Huang *et al*., 2017; Zhang *et al*., 2016b). Eleven imprinted genes related to seed size and 1,000-seed weight were obtained based on candidate gene-based association study. Among 11 imprinted genes, the gene *Lus10036044* (MEG) encoding a plant AT-rich sequence- and zinc-binding (PLATZ) transcription factor had significantly associated signal peaks on chromosome 1 (Table S8). PLATZ transcription factor is a novel class of plant-based zinc ion and DNA binding proteins, reported to regulate the seed size and weight (Azim *et al*., 2020). *ZmPLATZ12* (*Fl3*) is a maternally expressed imprinted gene specifically expressing in the starchy cells of endosperm in maize. The semi dominant negative *fl3* mutant resulted in severe defects of endosperm and dramatically reduced the weight of seeds (Li *et al*., 2017). In rice, the PLATZ transcription factor *GL6* positively controlled grain length through promoting cell proliferation in grains. The null *gl6* mutant led to short grains, whereas overexpression the *GL6* produced large grains (Wang *et al*., 2019). Another PLATZ gene *SHORT GRAIN6* (*SG6*) determined grain size by regulating the cell division of spikelet hull. The grain size and weight was significantly enlarged in the *SG6* overexpression lines and reduced in *sg6* mutant lines in rice (Zhou and Xue, 2020).

Another candidate gene *Lus10037040* (MEG) located on chromosome 1, which belongs to the MADS-box genes (Table S8). MADS-box genes had important functions in the development of seed by epigenetic mechanism including DNA methylation and histone modifications (Zhang *et al*., 2016b). In rice, *OsMADS87* (MEG) affected seed size by regulating endosperm cellularization during syncytial stage. Over expression the *OsMADS87* led to larger seeds, and *OsMADS87-*RNAi resulted in smaller seeds (Chen *et al*., 2016). The MADS-box gene *PHE1* (PEG) regulated seed size in *Arabidopsis thaliana* via influencing the expression of *AGL62* which might affect the endosperm cellularization (Sun *et al*., 2017). *OsMADS29* regulated seed development though regulating cell degeneration of maternal tissues. *OsMADS29*-RNAi resulted in aborted and/or shriveled seeds with deficient starch accumulation in endosperm (Yang *et al*., 2012). Heterologous expression the *CnMADS* gene significantly increased the seed size of *Arabidopsis* (Sun, 2018).

*Lus10024230* annotated as flavonol synthase (FLS) is also potentially involved in seed size control (Table S8). In the lines of *FLS*-RNAi of tobacco, the pods and seed development was arrested and the height, pods size, pods weight, seeds number were significantly reduced (Mahajan *et al*., 2011). Furthermore, the alternative alleles at *Lus10010350* (haplotypes B–K) had significantly longer seed length and width, and larger 1,000-seed weight than those in reference allele (haplotype A) in 200 flax accessions (Figure 7G-J, Figure S7). Together, our study identified a few candidate imprinted gene which are potentially involved in seed development and modulate the seed size. The genetic variation of these genes between flax lines may be harnessed as breeding tool for enhance seed yield.

### Intraspecific variation of flax imprinted genes

DNA methylation, histone modification and non-coding small RNAs caused genomic imprinting (Zhang *et al*., 2016a; Sha, 2008; Hanna and Kelsey, 2017). Epigenetic modification often varied across different individuals of the same species (Pignatta *et al*., 2014; Xu *et al*., 2019). In maize, differentially methylated regions (DMRs) were changed in different subgroups and genotypes (Xu *et al*., 2019; Li *et al*., 2015). In *Arabidopsis*, DNA methylation and small RNAs differentiated in natural populations and contributed to phenotypic diversity (Pignatta *et al*., 2014; Schmitz *et al*., 2011; Becker *et al*., 2011; Graaf *et al*., 2015; Schmitz *et al*., 2013). Genomic imprinting, as the functional product of epigenetic modification, varied within a same species and the intraspecific variation of imprinted genes was associated with epigenetic variation (Waters *et al*., 2013; Pignatta *et al*., 2014). In this research, the analysis of phylogenetic tree and PCA for imprinted SNPs showed that imprinted SNPs effectively divided the 200 flax germplasms into oil, OF and fiber flax subgroups (Figure 5, Figure S6), suggesting that genomic imprinting changed in different subgroups and contributed to phenotypic diversity in flax.

### Some imprinted genes show evidence of positive selection

According to previous reports, some imprinted genes showed positive selection features (Hutter *et al*., 2010; Berger *et al*., 2012). *MEA* as a component of FIS-PRC2 was a very important conserved imprinted gene in seed development underwent positive selection in the out-crossing lineages but not in the self-fertilizing species of *Arabidopsis* (Spillane *et al*., 2007; Miyake *et al*., 2009). Conserved imprinted genes displayed higher dN/dS rates than non-conservative imprinted genes between maize, rice and sorghum, suggesting conserved imprinted genes showing greater evidence of positive selection (Waters *et al*., 2013). Compared with MEGs, PEGs exhibited elevated dN/dS values and more likely to under positive darwinian selection in *Arabidopsis thaliana* (Tuteja *et al*., 2019). Our data showed that the nucleotide diversity of imprinted genes in oil flax subgroup was significantly higher than that in fiber flax subgroup (Figure 6A-C). The π values of some imprinted genes, such as *Lus10010350* (PEG, Figure 6D-F), *Lus10024230* (MEG, Figure 6G-I) and *Lus10041386* (MEG, Figure S8) were also significant difference between oil and fiber flax subgroup. Our results revealed that imprinted genes have been undergone artificial selection in the process of flax domestication from oil flax to fiber flax (Guo *et al*., 2020).

By analyzing the nucleotide diversity of imprinted genes in different flax subgroups, we found that the π values of imprinted genes in oil flax subgroup were significantly higher than those in fiber flax subgroup no matter what parental origin they were (Figure 6A-C). Meanwhile, we also discovered that the imprinted genes related to seed size and weight contained MEGs and PEGs (Table S8). It seemed that MEGs and PEGs were same shaped by selective force in flax population differentiation although the number of MEGs was larger than that of PEGs. So, we expected that imprinted genes undergo positive selection is related to the functions, but not to the parental origin which was different from the previous report (Tuteja *et al*., 2019). Compared with parental conflict theory, the imprinting under relaxed selection theory that genomic imprinting evolves consistent with neo-functionalization (Rodrigues and Zilberman, 2015) can better explain the intraspecific imprinting variation in flax subgroups.

## Materials and methods

### Plant Material and Tissue collection

The two parental lines of flax (*Linum usitatissimum* L.) for reciprocal crosses, CIli2719 (C) and Z11637 (Z), were grown at the Miquan Experiment filed in Urumqi, Xinjiang. The large seed line CIli2719 which 1000-seed weight was about 10.5g originated from France and the small seed line Z11637 that 1000-seed weight was about 3.7g originated from the United States. The seeds of CIli2719×Z11637 (CZ) and Z11637×CIli2719 (ZC) were collected at 7 DAP (day after pollination). Endosperm tissues were collected from at least 50 seeds by manual dissection in each replicate and were immediately frozen in liquid nitrogen. Three biological repeats were set up for each line. For phenotyping, the 200 accessions were planted in four environment comprising Dali in Yunnan Province in 2016 (2016DL), Urumqi in Xinjiang autonomous region in 2017 and 2019 (2017UR, 2019UR), and YiLi in Xinjiang autonomous region in 2019 (2019YL). Planting and phenotyping of the 200 accessions were performed using a same strategy as described in our previous study (Guo *et al*., 2020).

### Library construction for RNA-Seq

Total RNA was extracted using a RNAprep Pure Plant Kit (Tiangen Biotechnology of Beijing, http://www.tiangen.com/). The quantification and qualification of RNA was checked by 1% agarose gels, NanoPhotometer^®^ spectrophotometer (IMPLEN, CA, USA), Qubit^®^ RNA Assay Kit in Qubit^®^ 2.0 Flurometer (Life Technologies, CA, USA) and the RNA Nano 6000 Assay Kit of the Bioanalyzer 2100 system (Agilent Technologies, CA, USA). The RNA-seq libraries were generated using NEBNext® Ultra™ RNA Library Prep Kit for Illumina® (NEB, USA) according to the manufacturer’s instructions and the high-throughput sequencing was performed with the Illumina NovaSeq6000 platform. Then, the quality and quantity of these libraries were assessed by using the Agilent Bioanalyzer 2100 system and Q-PCR. A data size of 301.98 million 125bp paired-end raw reads was obtained from CZ and ZC.

### Read mapping and gene expression analysis

After removing the reads containing adapter, reads containing ploy-N (> 10%) and low quality reads (Q_phred_ ≤ 20) from raw data, a total of 288.37 million clean reads (43.26 Gb) were obtained for the following analysis. The clean reads were aligned to flax reference genome (https://phytozome.jgi.doe.gov/pz/portal.html#!info?alias=Org_Lusitatissimum) (Wang *et al*., 2012) using Hisat2 v2.0.4. HTSeq v0.9.1 to count the reads numbers mapped to each gene. And then the expected number of Fragments Per Kilo base of transcript sequence per Millions base pairs sequenced (FPKM) of each gene was calculated based on the length of the gene and reads count mapped to this gene (Trapnell *et al*., 2010). In the three biological replicates, the gene with an average expression level of FPKM > 1 was identified as “expressed” (Meng *et al*., 2018).

### Identification of imprinted genes

The clean reads from CZ and ZC were aligned to parental genomes CIli2719 and Z11637 from our previous research (Guo *et al*., 2020) to obtain the reads of C and Z alleles at each SNP site for parental allelic expression analysis. Theoretically, the allelic ratio of the maternal to paternal is 2 to 1 in hybrid endosperm. Based on the 2m:1p ratio, SNP loci with more than 10 alleles reads in reciprocal crosses were used to perform a two-tailed chi square (χ^2^) test. Moderately imprinted SNP loci had significant allelic bias (χ^2^<0.05) and >80% of the transcripts from the maternal allele for maternally expressed imprinted SNP loci or >60% of the transcripts coming from the paternal allele for paternally expressed imprinted SNP loci in both reciprocal hybrids. Strong maternally or paternally expressed imprinted SNP loci were defined as having significant allelic bias (χ^2^<0.01) and >90% of transcripts derived from the maternal allele or paternal allele, respectively. Complete maternally/paternally expressed imprinted SNP loci had >99% of the transcripts from the maternal/paternal allele (Waters *et al*., 2013; Meng *et al*., 2018). And the genes containing at least one imprinted SNP loci were identified as imprinted genes.

### Validation of imprinted gene and expression analysis

Thirteen genes were used to perform Quantitative RT-PCR (qRT-PCR) analysis (Table S9) and twelve imprinted genes were detected using a PCR-sequencing method (Table S10) (Meng *et al*., 2018; Xu *et al*., 2014). The endosperm cDNA samples at 7-DAP were collected for RNA isolation with three biological repeats for each sample.

The extraction, quantification and identification of total RNA were the same as that of library construction for RNA-Seq. First-strand cDNA synthesis was performed using 5×All-In-One RT MasterMix (with AccuRT Genomic DNA Removal Kit) according to the manufacturer recommended protocol for qRT-PCR and RT-PCR (abm, Cat. No.G492, http://www.abmGood.com/). Each qRT-PCR reaction of CZ and ZC was performed by the manufacturer’s instructions of EvaGreen Express 2×qPCR MasterMix (abm, Cat. No.MasterMix-ES, http://www.abmGood.com/) and BioRad^®^CFX96 Real-Time PCR system (Bio-Rad). Relative expression was quantified with the geometric mean of internal reference genes *ETIFI* (eukaryotic translation initiationfactor 1), *GAPDH* (glyceraldehyde 3-phosphate dehydrogenase), and *ETIF5A* (eukaryotic translation initiationfactor 5A) (Hobson and Deyholos, 2013; Huis *et al*., 2010). For RT-PCR, a 400-800bp amplification fragment of each gene was amplified by different primers with four endosperm cDNA samples: CC (inbred lines of CIli2719), ZZ (inbred lines of Z11637), CZ and ZC of 7-DAP endosperm. The RT-PCR amplified products contained at least one imprinted SNP sites were analyzed on agarose gels and then sequenced.

### Functional characterization of imprinted features

Gene annotation of imprinted genes in flax endosperm was downloaded from the reference genome (https://phytozome.jgi.doe.gov/pz/portal.html), and GO enrichment analysis was carried out with WEGO (http://wego.genomics.org.cn/) (Xu *et al*., 2014; Ye *et al*., 2006).

The imprinted genes of flax were investigated for sequence homology in *Arabidopsis* (Gehring *et al*., 2011), rice (Yuan *et al*., 2017; Luo *et al*., 2011), maize (Waters *et al*., 2013; Dong, 2017), sorghum (Zhang *et al*., 2016a),and castor bean (Xu *et al*., 2014) using blast. The peptide sequences of flax imprinted genes were obtained from the flax database in Phytozome v12.1 (https://phytozome.jgi.doe.gov/pz/portal.html). Then, the peptide sequences were aligned to the *Arabidopsis* genome (*Arabidopsis thaliana* TAIR10) and high-scoring (E-value<1E-10, 1E-20 and 1E-50) blasts hits were ordered by increasing E-value. If the *Arabidopsis* imprinted genes were identified amongst the blast hits, and the gene with the smallest E-value was recorded. Similarly, candidate genes from this study in flax were aligned to the rice (*Oryza sativa* v7_JGI), maize (*Zea mays* Ensembl-18), sorghum (*Sorghum bicolor* v3.1.1) and castor bean genome (*Ricinus communis* v0.1). The Venn diagrams were drawn by the draw venn diagram online software (http://bioinformatics.psb.ugent.be/webtools/Venn/).

For clustering analysis of imprinted genes, a standard was applied that imprinted genes within 10 kb of one another in the flax genome was a candidate cluster (Gehring *et al*., 2011; Xu *et al*., 2014). Positions of imprinted genes on chromosome were mapped using the MapChart software (Voorrips, 2002).

### The tissue-specific expression analysis of imprinted genes in endosperm

The gene-expression patterns for MEGs and PEGs in various flax tissues in reciprocal hybrids were identified based on RNA-seq analysis. The endosperm and embryo tissues were harvested at 7 DAP and the leaf tissues were collected at 2 weeks after planting. For each sample, three biological replicates were used. The FPKM expression values of all genes and imprinted genes in CZ and ZC were log-transformed. All genes with FPKM>1 in endosperm were used in this study. The heat map and hierarchical clustering of normalized expression levels (FPKM) were performed with the MeV4.9.0 software (Multi Experiment Viewer, https://sourceforge.net/projects/mevtm4/files/mev-tm4/) (Guo *et al*., 2020).

### Phylogenetic tree and population structure analysis using imprinted SNPs

To test the relationship between SNP variation at population level and population structure, an individual-based neighbor-joining tree was generated based on the all imprinted SNPs (498 SNPs), ME-SNPs (319 ME-SNPs) or PE-SNPs (179 PE-SNPs) by TASSEL’s Cladogram function (Bradbury *et al*., 2007). We compared the kinship of 200 accessions calculated by imprinted SNPs (498) and genome-wide SNPs (674,074) obtained from our previous research (Guo *et al*., 2020). Principal component analysis (PCA) was conducted based on the imprinted SNPs by using the software TASSEL 5.0 (Bradbury *et al*., 2007).

### Identifying selection signatures of imprinted genes

A set of 78 oil flax and 51 fiber flax accessions which represent two primary morphotypes of cultivated flax were used for selective sweeps analysis. To test the genetic diversity of imprinted genes in different subgroups, 3,191 SNPs were obtained by mapping the imprinted genes to our previously constructed variation map (Guo *et al*., 2020). The π values of all imprinted genes (241), MEGs (143) and PEGs (98) between oil and fiber subgroups were calculated at the gene level using all SNPs within each imprinted gene by the software DnaSP 5.1 (Librado and Rozas, 2009). Furthermore, the π values were also calculated at the SNP level in oil subgroup and fiber subgroup for detecting the selection signatures in a single imprinted gene using DnaSP 5.1 (Librado and Rozas, 2009).

### Candidate gene-based association study for seed size-related traits using flax imprinted genes

To analyze the association between imprinted genes and seed size-related traits, imprinted genes were used to perform candidate gene-based association study by the general linear model (GLM) and mixed linear model (MLM) in TASSEL 5.0 (Bradbury *et al*., 2007). The 3,191 SNPs in the imprinted genes and seed size-related traits including seed length (SL), seed width (SW) and 1,000-seed weight (1,000-SW) of 200 flax accessions were obtained from our previous research (Guo *et al*., 2020). For GLM analysis, the top two principal components (PC) were used to generate the population structure matrix and the threshold was set as 0.1/total SNPs (log_10_(*P*) =-4.50). For MLM analysis, P matrix and Kinship (K) matrix need to be considered, and the suggestive threshold was set as 1/total SNPs (log_10_(*P*) = -3.50). The imprinted genes repeatedly detected for at least two environments or traits were considered to be associated with seed size and weight.

## Acknowledgements

We thank Chinese Crop Germplasm Resources Information System (CGRIS), Plant Gene Resource Centre in Canada (PGRC) and United States National Plant Germplasm System (U.S.NPGS) for providing us with germplasm resources. This study was supported by the National Natural Science Foundation of China (32060426), Resource Platform Project of Xinjiang Uygur Autonomous Region of China (PT1808) and Science and Technology Innovation Project for Doctoral Students of Xinjiang University of China (XJUBSCX-2017017). We are grateful to professors Ming Luo and Donghe Xu for constructive comments to improve the manuscript. The authors declare no competing interests of this work.

## Availability of data and materials

Raw resequencing sequence data of the two parental lines CIli2719 and Z11637 of flax for reciprocal crosses are available at NCBI under accession PRJNA590636 (ncbi.nlm.nih.gov/bioproject/PRJNA590636) (Guo *et al*., 2020).

## Supplementary information

**Figure S1**. Flow chart for identification of imprinted genes in flax endosperm.

**Figure S2**. Verification of thirteen genes in flax endosperm based on qRT-PCR analysis. Thirteen genes were chosen for the qRT-PCR analyses. Among these genes, five were MEGs, five were PEGs, and others were not imprinted genes. So, the gene expression level between qRT-PCR and RNA sequencing of these thirteen genes represented the whole types of genes in this study.

**Figure S3**. Validation of the imprinted genes in flax endosperm by PCR sequencing. Twelve imprinted genes including nine MEGs and three PEGs were selected for validation. Each gene was designed by a pair of primers with a 400-800bp amplification fragment which was a part of the corresponding CDS sequence of CC (endosperm of CIli2719 self-cross), ZZ (endosperm of Z11637 self-cross), CZ (endosperm of CIli2719×Z11637), and ZC (endosperm of Z11637×CIli2719) and the amplification fragment contained at least one imprinted SNP site.

**Figure S4**. Gene ontology analysis of 229 identified imprinted genes and 115 conserved imprinted genes. (A) Gene ontology analysis of identified imprinted genes. MEGs represented 135 maternally expressed genes (red), PEGs represented 94 paternally expressed genes (purple), All imprinted genes represented 229 moderate imprinted genes (blue), All genes represented all endosperm-expressed genes with at least ten reads could be assigned to a specific allele in both CZ and ZC (yellow). (B) Gene ontology analysis of conserved imprinted genes. Conserved MEGs represented 75 conserved maternally expressed genes (red), Conserved PEGs represented 40 conserved paternally expressed genes (purple), Conserved imprinted genes represented 115 conserved imprinted genes (blue), All genes were presented as A.

**Figure S5**. Expression of conserved imprinted genes in different tissues of flax in both reciprocal hybrids CZ and ZC based on RNA-seq analysis. (A, B) The gene-expression patterns for MEGs (A) and PEGs (B) of 115 conserved imprinted genes. (C) The gene-expression patterns for MEGs and PEGs of 38 conserved strong imprinted genes. endo-MEGs, MEGs that expressed preferentially in endosperm; con-MEGs, MEGs that expressed in many tissues; endo-PEGs, PEGs that expressed preferentially in endosperm; con-PEGs, PEGs that expressed in many tissues. The normalized values were used for hierarchical clustering and the heat map indicates relative levels of expression. The endosperm and embryo tissues were harvested at 7 DAP and the leaf tissues were collected at 2 weeks after planting. For each sample, three biological replicates were used.

**Figure S6**. MEGs and PEGs can differentiate flax subgroups. (A) Phylogenetic tree of 200 flax accessions inferred from ME-SNPs. (B) Phylogenetic tree of 200 flax accessions inferred from PE-SNPs. ME-SNPs, maternally expressed imprinted SNP loci; PE-SNPs, paternally expressed imprinted SNP loci. Fiber flax, oil-fiber dual purpose flax (OF), and Oil flax were represented in red, blue and green colors, respectively.

**Figure S7**. Boxplots for seed length and seed width based on the haplotypes (Hap.) for *Lus10010350*. (A-B) The seed length (A) and seed width (B) in 2016DL. (C-D) The seed length (C) and seed width (D) in 2017UR. (E-F) The seed length (E) and seed width (F) in 2019UR. (G-H) The seed length (G) and seed width (H) in 2019YL. In the box plots, the center line represented the median, box limits indicated the upper and lower quartiles, whiskers marked the range of the data and points showed outliers. *n* indicates the number of accessions with the same genotype. The difference between haplotypes was analyzed by two-tailed *t* tests.

**Figure S8**. The nucleotide diversity distribution of *Lus10041386*. (A) The nucleotide diversity distribution of *Lus10041386* on chromosome 15 among Oil and Fiber subgroups. (B) Boxplots for nucleotide diversity of *Lus10041386* among Oil and Fiber subgroups. The difference was analyzed by two-tailed *t* tests.

**Table S1**. Imprinted genes in both hybrid endosperms and associated SNPs.

**Table S2**. Functional annotations of 229 imprinted protein-coding genes.

**Table S3**. Gene ontology enrichment analysis of 229 flax imprinted genes.

**Table S4**. The conservation of imprinted genes detected in our endosperm samples.

**Table S5**. The number of conserved flax imprinted genes with other species.

**Table S6**. Gene ontology enrichment analysis of 115 conserved imprinted genes in flax.

**Table S7**. Clusters of imprinted genes.

**Table S8**. Imprinted genes related to seed size and 1000-seed weight based on candidate gene-based association study.

**Table S9**. Primers for qRT-PCR.

**Table S10**. Primers for imprinting validation by PCR-sequencing.

