## Supplemental Figure S1-S8 and Table S1-S10 for "Genome-wide screen of genomic imprinting in endosperm and population-level analysis reveal allelic variation for imprinting in flax": Supplemental Figure S1-S8.docx

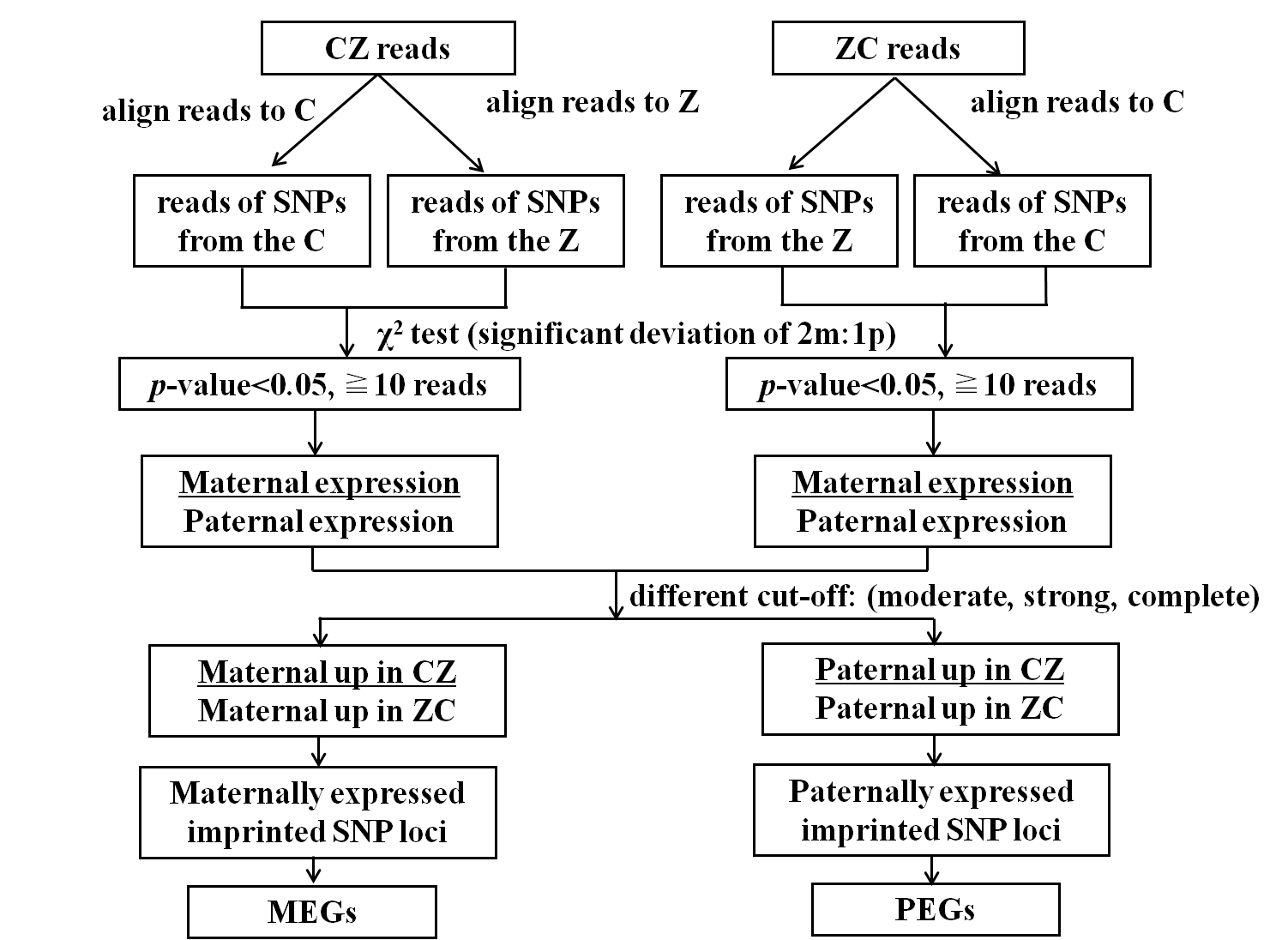

**Figure S1.** Flow chart for identification of imprinted genes in flax endosperm.

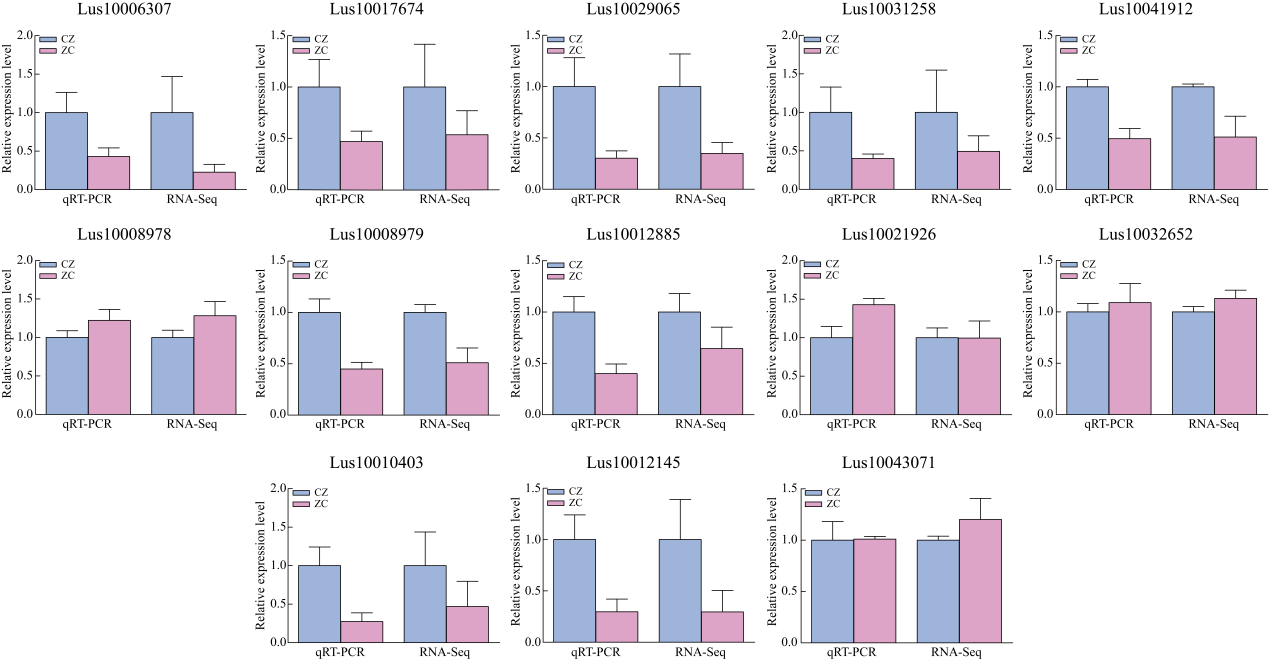

**Figure S2.** Verification of imprinted genes in flax endosperm based on qRT-PCR analysis. Thirteen genes were chosen for the qRT-PCR analyses. Among these genes, five were MEGs, five were PEGs, and the others were non-imprinted genes. So, the gene expression level between qRT-PCR and RNA sequencing of these thirteen genes represented the whole types of genes in this study.

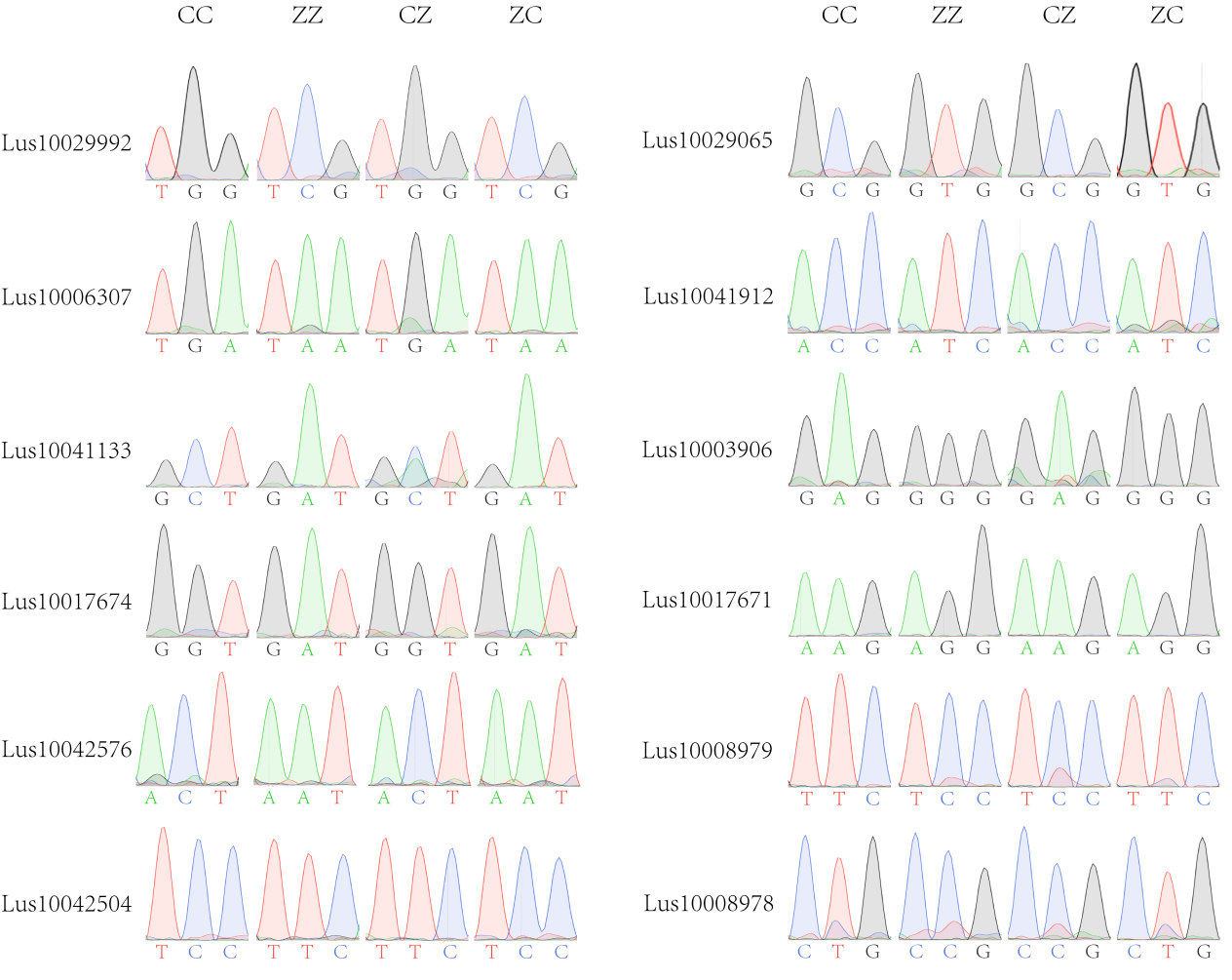

**Figure S3.** Validation of the imprinted genes in flax endosperm by PCR sequencing. Twelve imprinted genes including nine MEGs and three PEGs were selected for validation. Each gene was designed by a pair of primers with a 400-800bp amplification fragment which was a part of the corresponding CDS sequence of CC (endosperm of CIli2719 self-cross), ZZ (endosperm of Z11637 self-cross), CZ (endosperm of CIli2719×Z11637), and ZC (endosperm of Z11637×CIli2719) and the amplification fragment contained at least one imprinted SNP site.

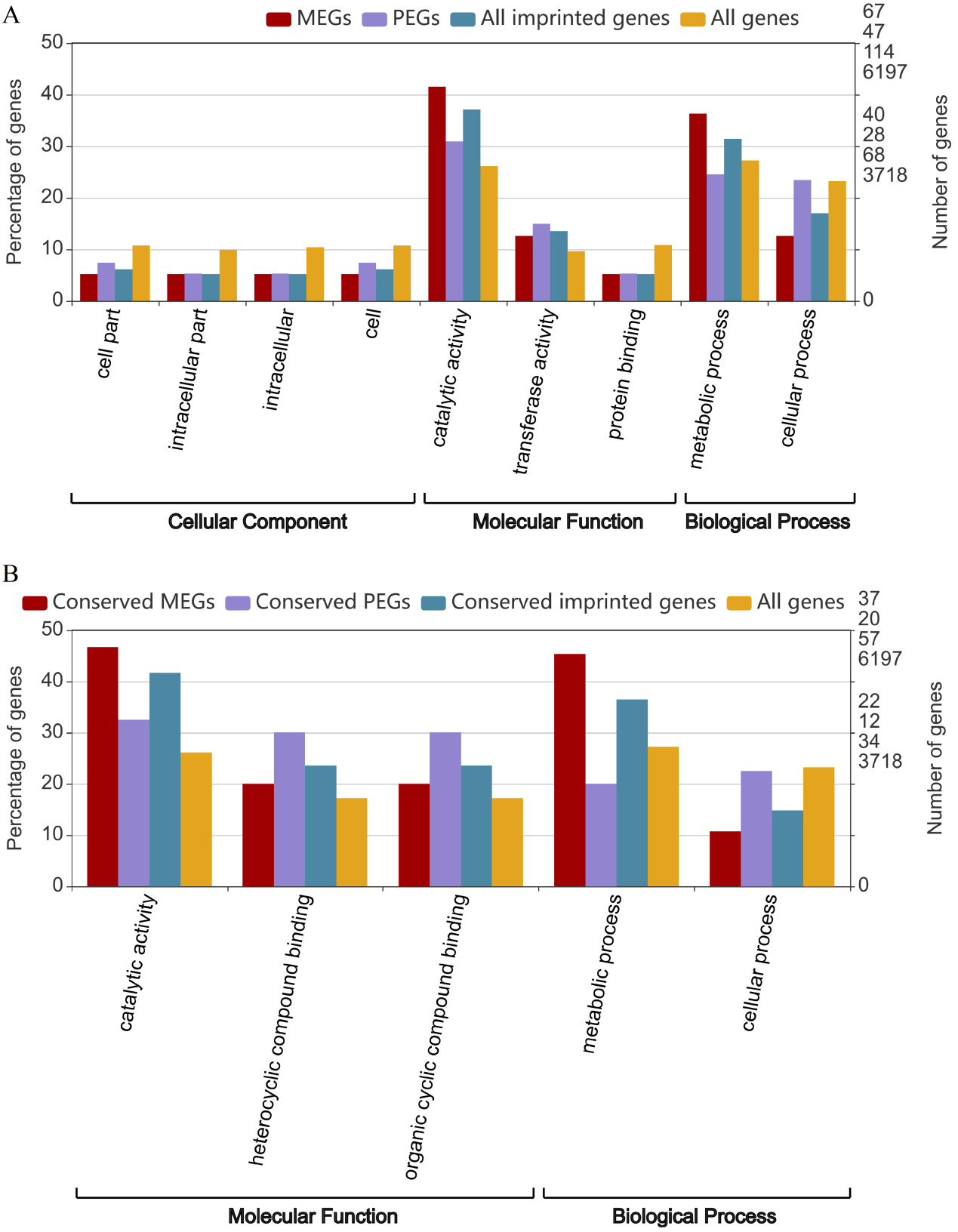

**Figure S4.** Gene ontology analysis of 229 protein-coding imprinted genes and 115 conserved imprinted genes. (A) Gene ontology analysis of 229 protein-coding imprinted genes. MEGs represented 135 maternally expressed genes (red), PEGs represented 94 paternally expressed genes (purple), All imprinted genes represented 229 moderate imprinted genes (blue), All genes represented all endosperm-expressed genes with at least ten reads could be assigned to a specific allele in both CZ and ZC (yellow). (B) Gene ontology analysis of conserved imprinted genes. Conserved MEGs represented 75 conserved maternally expressed genes (red), Conserved PEGs represented 40 conserved paternally expressed genes (purple), Conserved imprinted genes represented 115 conserved imprinted genes (blue), All genes were presented as A.

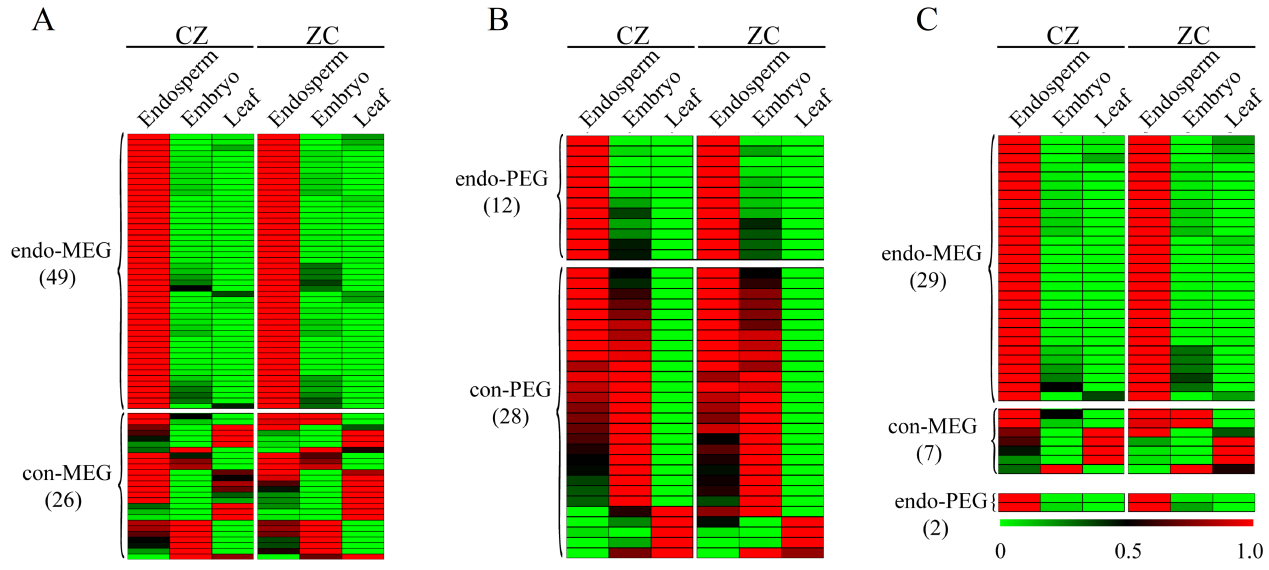

**Figure S5.** Expression of conserved imprinted genes in different tissues of flax in both reciprocal hybrids CZ and ZC based on RNA-seq analysis. (A-B) The gene-expression patterns for MEGs (A) and PEGs (B) of 115 conserved imprinted genes. (C) The gene-expression patterns for MEGs and PEGs of 38 conserved strong imprinted genes. endo-MEGs, MEGs that expressed preferentially in endosperm; con-MEGs, MEGs that also expressed in other tissues; endo-PEGs, PEGs that expressed preferentially in endosperm; con-PEGs, PEGs that also expressed in other tissues. The normalized values were used for hierarchical clustering and the heat map indicates relative levels of expression. The endosperm and embryo tissues were harvested at 7 DAP and the leaf tissues were collected at 2 weeks after planting. For each sample, three biological replicates were used.

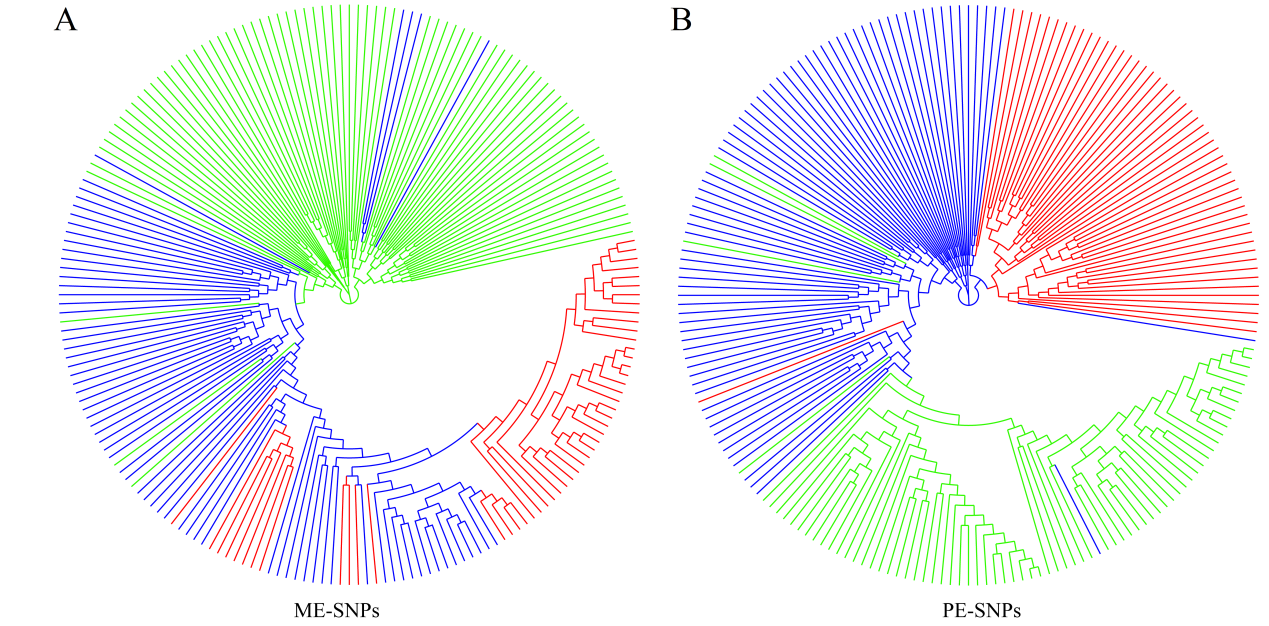

**Figure S6.** MEGs and PEGs can differentiate flax subgroups. (A) Phylogenetic tree of 200 flax accessions inferred from ME-SNPs. (B) Phylogenetic tree of 200 flax accessions inferred from PE-SNPs. ME-SNPs, maternally expressed imprinted SNP loci; PE-SNPs, paternally expressed imprinted SNP loci. Fiber flax, oil-fiber dual purpose flax (OF), and Oil flax were represented in red, blue and green colors, respectively.

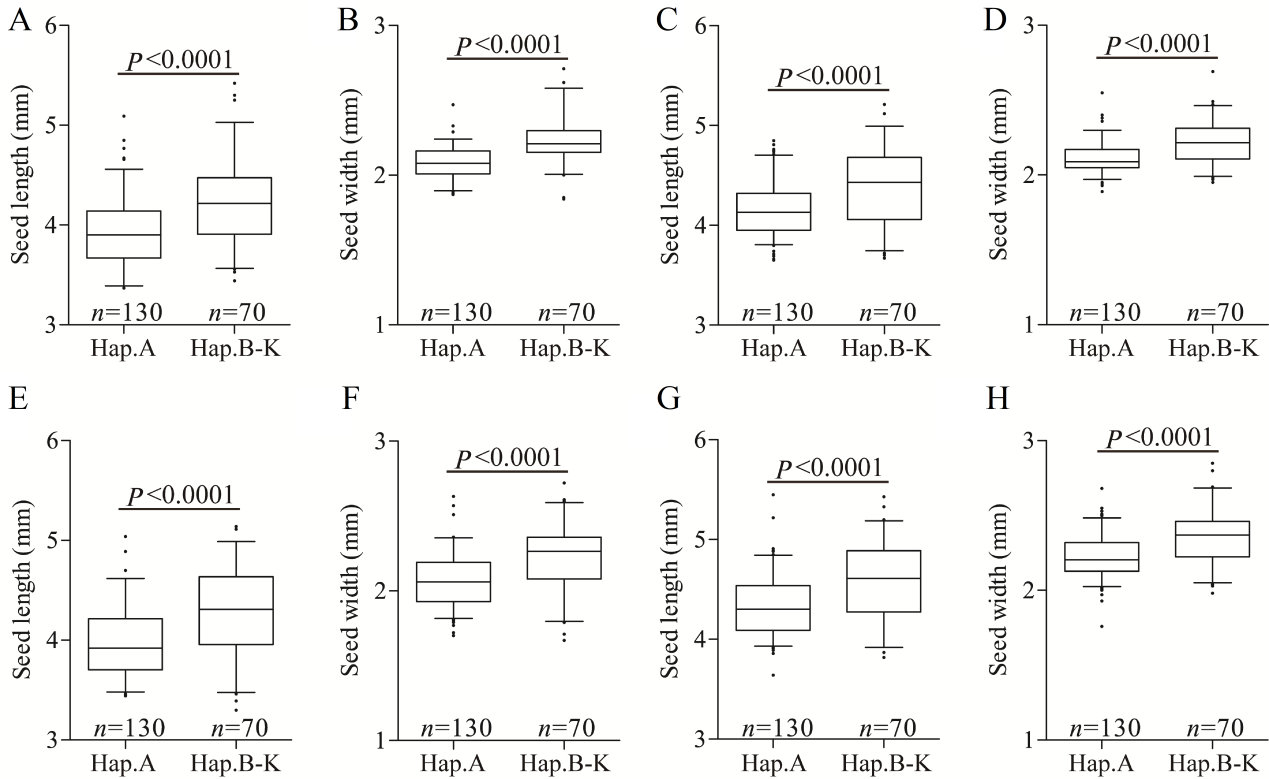

**Figure S7.** Boxplots for seed length and seed width based on the haplotypes (Hap.) for *Lus10010350.* (A-B) The seed length (A) and seed width (B) in 2016DL. (C-D) The seed length (C) and seed width (D) in 2017UR. (E-F) The seed length (E) and seed width (F) in 2019UR. (G-H) The seed length (G) and seed width (H) in 2019YL. In the box plots, the center line represented the median, box limits indicated the upper and lower quartiles, whiskers marked the range of the data and points showed outliers. *n* indicates the number of accessions with the same genotype. The difference between haplotypes was analyzed by two-tailed *t* tests.

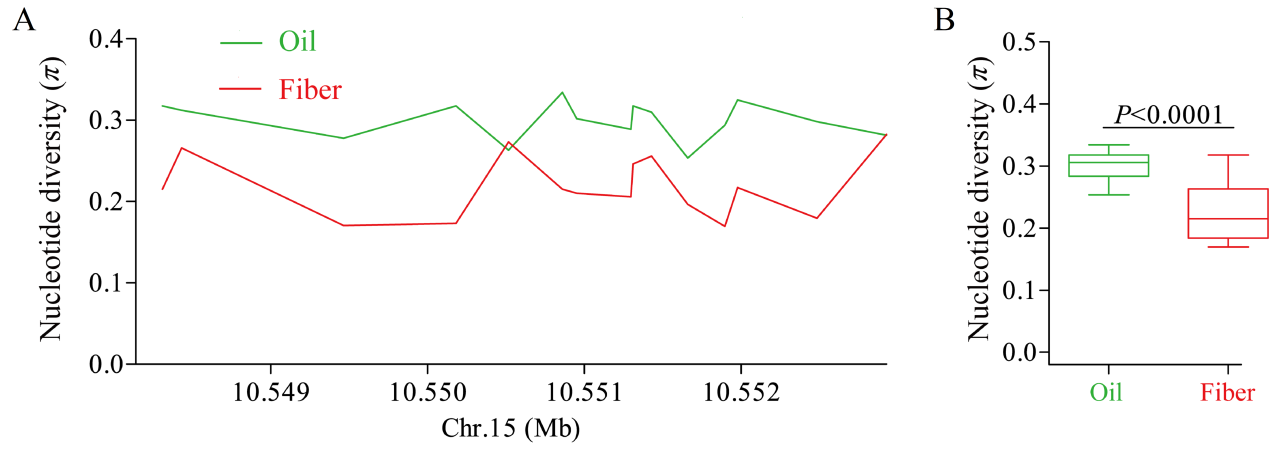

**Figure S8.** The nucleotide diversity distribution of *Lus10041386*. (A) The nucleotide diversity distribution of *Lus10041386* on chromosome 15 among Oil and Fiber subgroups. (B) Boxplots for nucleotide diversity of *Lus10041386* among Oil and Fiber subgroups. The difference was analyzed by two-tailed *t* tests.
